# Metabolic rewiring of cancer cells induces metastasis via ERK5 but triggers recognition by NK cells

**DOI:** 10.1101/2025.08.26.672105

**Authors:** Sara Zemiti, Steffy Escayg, Benjamin Hay, Amel Bouzid, Catherine Alexia, Carine Jacquard, Noemie Dudzinska, Ananya Sen, Jose Prudenciano, Daouda Abba Moussa, Michael Constantinides, Loïs Coenon, Nerea Allende-Vega, Mauricio Campos Mora, Sabine Gerbal-Chaloin, Alberto Anel, Marion De Toledo, Guillaume Bossis, Ana Bellia Aznar, Susana Prieto, Daniel Fisher, Farida Djouad, Yoan Arribat, Jean-François Rossi, Martin Villalba, Delphine Gitenay

## Abstract

Metastasis is largely controlled by Natural Killer (NK) cell-mediated immune surveillance. To colonize new environments, cancer cells undergo epithelial to mesenchymal transition (EMT), which allows them to detach and migrate. EMT is fueled by fatty acid oxidation (FAO), which partially replaces glycolytic-based tumor metabolism. Whether metabolic rewiring affects the targeting of cancer cells by NK cells remains unknown. Here, we show that forcing solid cancer cells to perform FAO by inhibiting pyruvate dehydrogenase kinase 1 with dichloroacetate (DCA) activates extracellular signal-regulated kinase-5 (ERK5), triggering EMT and tumor cell migration and invasion. Concomitantly, FAO induces the expression of ligands that mediate NK cell recognition. Consequently, NK cells better infiltrated DCA-treated 3D tumor spheroids, where they exerted their cytotoxic effects. DCA-treated cells showed increased migration in a zebrafish model, whereas metastasis from mammary cancer cells grafted into immune-deficient mice was enhanced by DCA. These migrating/metastatic cells are preferentially killed by NK cells, which strongly limit their invasive potential. Hence, FAO promotes both metastasis and NK-mediated tumor surveillance, highlighting the Achilles’ heel of metastatic cells, which may offer new therapeutic opportunities.

**Teaser:** Metastasis recognition and killing by immune cells, such as NK cells, requires a metabolic shift that relies on lipid metabolism.

## Introduction

Metabolism is altered in cancer cells, which rely on glycolytic pathways to meet the high-energy demands for growth and tumor maintenance. Thus, aerobic glycolysis is a hallmark of cancer cells(*1*). However, other metabolic changes have also been identified, such as modifications in amino acid or fatty acid (FA) metabolism. Some of these lead to various metabolic alterations, resulting in simultaneous increases in mitochondrial respiration and glycolysis(*2*). Additionally, cancer cells adapt to changing environments by adjusting their metabolism, such as when they colonize new tissues and acclimatize to these conditions. This migration, termed metastatic colonization, is highly inefficient, because metastatic cells must overcome numerous obstacles to establish new tissues(*3*). Once metastatic lesions form, current treatments often fail to provide a lasting response. During their journey to new target tissues, tumor cells face a challenging metabolic environment and respond by increasing FA consumption and energy production via FA β-oxidation (FAO)(*4*). Therefore, lipid oxidation, which fuels β-oxidation and mitochondrial respiration, is linked to tumor progression, invasion, and the metastasis. The lipid transporter CD36 is considered a marker of metastasis-initiating cells(*5, 6*) and plays an important role in epithelial-to-mesenchymal transition (EMT) and tumor cell migration(*6, 7*).

Acyl-CoA synthetases (ACSL) catalyze the conversion of long-chain FAs to acyl-CoA, which is critical for phospholipid and triglyceride synthesis, lipid modification of proteins, and FAO(*8*). ACSL enzymes play a role in EMT (*9, 10*), which is an essential step for cancer cell detachment from the primary tumor to initiate metastasis. During migration through the host, metastatic cells are exposed to immune cells, particularly cytotoxic lymphocytes, which patrol blood and lymph. For example, natural killer (NK) cells mediate metastasis-specific I mmunosurveillance(*11*), and their number and activity are correlated with the quantity of circulating tumor cells and metastasis in several cancers(*12*). However the mechanisms by which NK cells recognize and kill metastatic cells remain unknown. In a model of TGFβ-induced EMT, Chockley *et al*. showed that E-cadherin (E-Cad) is recognized by the killer cell lectin like receptor G1 (KLRG1), an inhibiting receptor in NK cells(*11*). Cell adhesion molecule 1 (CADM1) is recognized by the cytotoxic and regulatory T cell molecule (CRTAM), an NK cell-activating receptor(*11*). Thus, decreased E-cadherin and increased CADM1expression enhanced metastatic cell recognition by NK cells. This may partially explain how metastatic cells are recognized, although merely losing the inhibitory E-Cad/KLRG1 interaction is insufficient to restore NK cell cytotoxicity(*11*). NK cells also control dormant tumor cells that can give rise to cancers, long after the primary tumors have been eliminated(*13*).

NK cells recognize targets by identifying their surface ligand patterns. Several ligands of NK cell-activating receptors, such as Poliovirus Receptor (PVR, also known as CD155), UL16 binding protein 2 (ULBP2), ULBP4, and MHC class I polypeptide-related sequences A/B (MICA/B), increase during EMT (*14*). The activating natural killer group 2D (NKG2D) receptor binds to MICA/B and various ULBPs, leading to target cell killing(*15*). Conversely, major histocompatibility complex-I (MHC-I), an inhibitory ligand(*11, 16*), decreases during EMT(*17*). Therefore, multiple mechanisms during EMT may result in the expression of NK cell-recognized ligands, supporting immune surveillance and partially explaining the inefficacy of metastatic cells in colonizing new environments(*12, 18*).

Metabolic changes significantly regulate cancer cell recognition by the immune system by altering the activation/inhibition balance of ligands on the tumor cell surface(*19*). For instance, forcing cells to perform OXPHOS induces MHC-I expression in leukemic cells (*19–22*); however, MHC-I is thought to decrease in metastatic cells (*23–25*), which partially explains their sensitivity to NK cells. In addition, metformin, which reduces glycolysis, induces Intercellular Adhesion Molecule-1 (ICAM-1), enhancing sensitivity to NK cell-mediated cell death(*26–28*). Similarly, the pyruvate dehydrogenase kinase-1 (PDK1) inhibitor dichloroacetate (DCA), which activates pyruvate dehydrogenase (PDH) to allow pyruvate entry into the Krebs cycle(*29, 30*), sensitizes leukemic cells to NK cells (*27*). In hematological cancer cells, DCA induces the expression of several genes involved in lipid metabolism such as the low density lipoprotein receptor (LDLR), which is a key regulatory receptor in cholesterol metabolism, CD36 and ACADVL through the extracellular-regulated protein kinase 5 (ERK5) pathway(*31, 32*). ERK5 also regulates immune cell recognition(*20, 21, 33*) and plays multiple roles in cancer hallmarks(*34*), including increased cell motility(*34, 35*) and a central role in EMT(*34, 36*). ERK5 thus appears as a promising candidate for linking metabolic alterations, EMT, and immune surveillance.

We hypothesized that increased lipid consumption by cancer cells could induce EMT and initiate the early stages of metastasis, thereby generating activating signals for NK cells. This metabolic stress might also induce stress ligands that activate NK cells, promoting tumor recognition and killing.

## Results

### DCA increases lipid metabolism, FAO and respiration

To develop a cell model with increased FAO, HCT116 cells were treated for 48h with 5mM DCA alone or in combination with 3µM of the FA oleic acid (OA). DCA increased neutral lipid content, as shown by confocal microscopy (Fig. 1A) and flow cytometry (Fig. 1B), even more in the presence of OA. As in leukemic cells(*31, 32*), DCA increased *ACADVL* and *CD36* mRNA levels in HCT116 cells (Figs. 1C and D). Transfection with siRNA against *ERK5* (siERK5) partially reversed the effects of DCA on *ACDVL*, but reversed totally those on*CD36* (Fig. 1C and D). DCA also increased CD36 expression at the plasma membrane (Fig. 1E). Adding OA or DCA to the cell culture medium was sufficient to increase the oxygen consumption (Fig. 1F).

**Figure 1.**
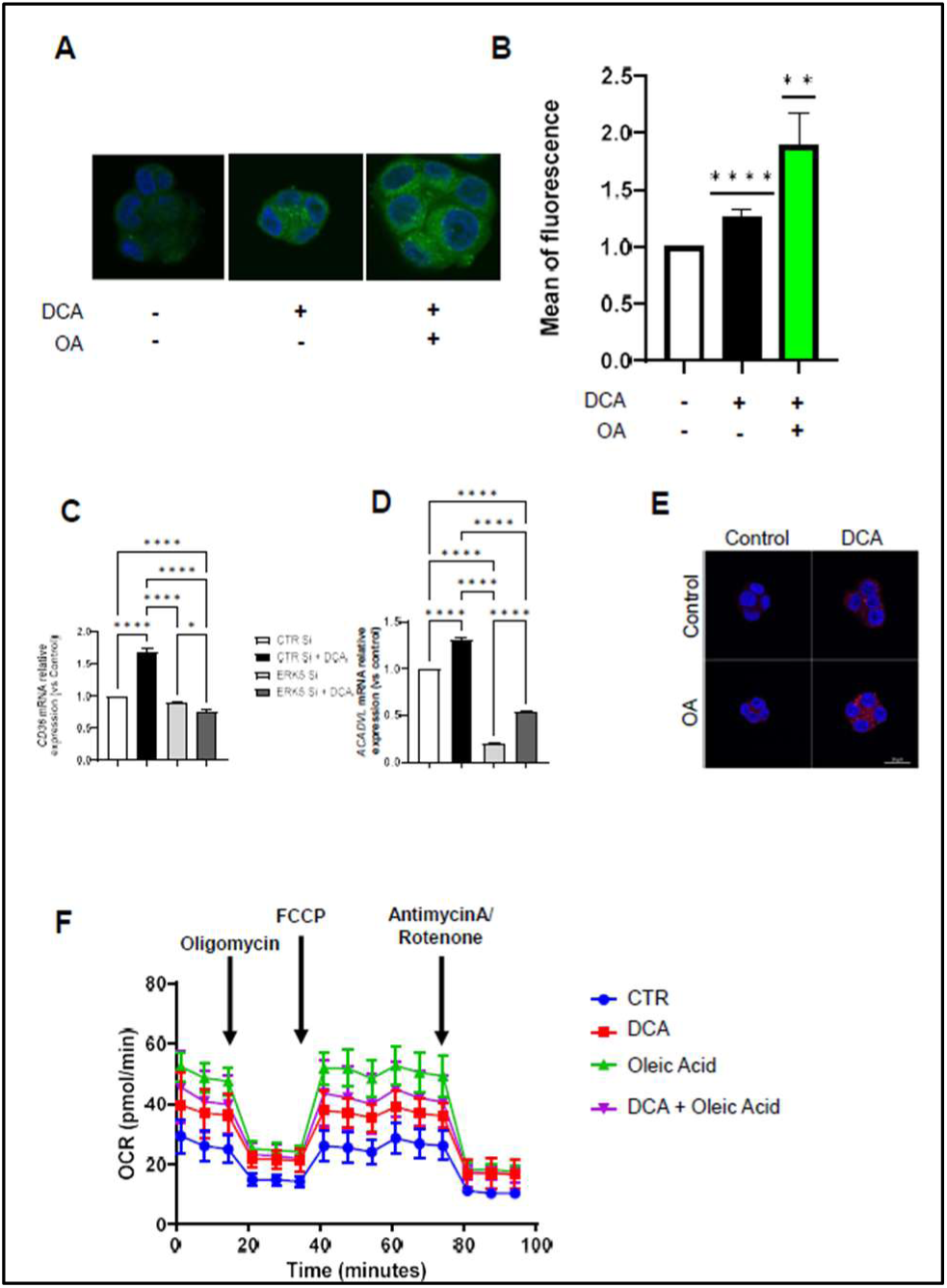
DCA increases lipid metabolism and oxidative phosphorylation (OXPHOS) in HCT116 cells. A-B) HCT116 cells were treated with 5mM DCA and 3µM oleic acid (OA) for 48h. Cells were incubated with Bodipy and DAPI and analyzed by immunofluorescence (A) or by flow cytometry (B) in 3 independent experiments. C-D) Cells were transfected with a siRNA control (siControl) or with a siRNA for ERK5 (siERK5). Forty-eight hours after transfection 200000 cells/well were seeded in 6 wells plates and treated with 10mM DCA. After 48h, RNA was extracted and the relative mRNA level of *ACADVL* (C) and *CD36* (D) were assessed and normalized to *β-actin* mRNA (n=3). E) HCT116 were treated with 5mM DCA and/or 3µM OA. After 48h, CD36 expression (in red) at the cell surface was assessed by microscopy (n=3). F) DCA treatment increases basal and maximal respiration in HCT116 cells. Cells were treated as in (A) and the oxygen consumption rate (OCR) was analyzed in a Seahorse. Statistical analysis was performed using one-way Anova followed by a Tukey multiple comparison test. Graphs represent mean +/- SD. *\* p < 0*.*05, ** p < 0*.*01, *** p < 0*.*001, **** p < 0*.*0001*; Tukey’s test was used to compare condition to control or as depicted in the graphic.

### DCA-induced lipid metabolism causes expression of EMT-related genes and a migratory phenotype

Because DCA induces lipid metabolism, we hypothesized that it could favor a metastatic phenotype. DCA increased the mRNA levels of several genes critical for EMT, *ZEB1, SNAIL*, and *TWIST* (Fig. 2A). siERK5 partially reversed the effects of DCA on *ZEB1* and *SNAIL* and in total on *TWIST* (Fig. 2A). We confirmed the induction of SNAIL at the protein level, whereas the ERK5 inhibitor XMD8-92 reversed DCA effects (Fig. 2B). Hence, ERK5 mediates DCA effects on EMT.

**Figure 2.**
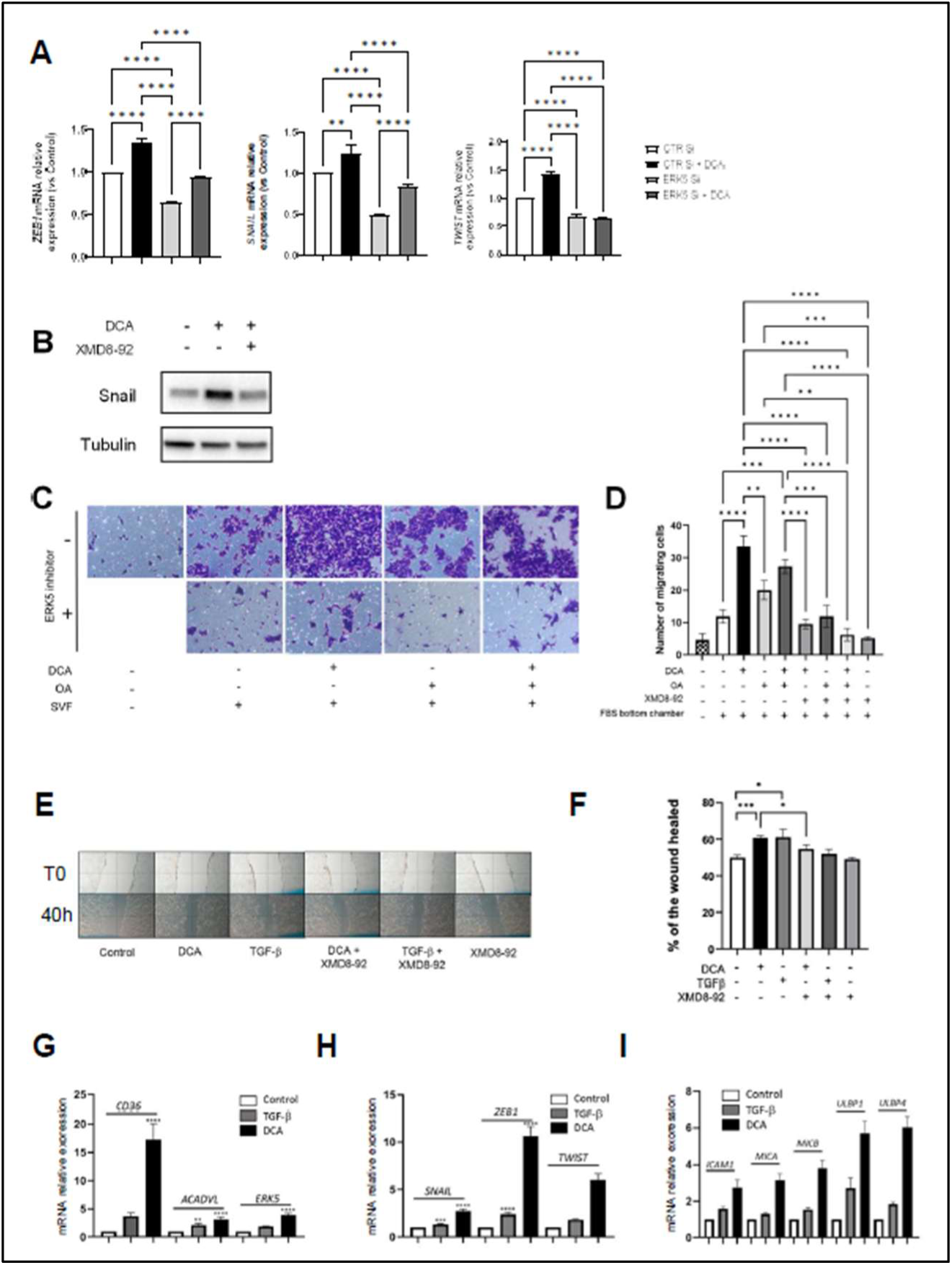
FAO induces EMT and a migratory phenotype in 2-Dimension (2D) through the ERK5 pathway. A) Twenty-four hours after seeding and at 60% confluency, HCT116 cells were transfected with either siControl or siERK5 and 48h later treated with 10mM DCA. After 48h, RNA was extracted and the relative mRNA level of *ZEB1, SNAIL* and *TWIST* were assessed and normalized to *β-actin* mRNA levels (n=3). B) Cells were seeded and treated for 48h with 20mM DCA and 5 µM of the ERK5 inhibitor XMD8-92. SNAIL and tubulin expression were analyzed by western blot. C) 7500 HCT116 cells were seeded in inserts of a transwell system in a FBS-deprived medium and incubated with 10mM DCA, 3µM OA and/or 5µM of XMD8-92. The lower part of the transwell contained 10% FBS to attract cells. After 48h we analyzed the number of colonies formed on the other side of the insert after fixation and crystal violet staining (representative of 3 experiments). D) On the pictures from C), 3 fields of the 3 independent experiments were analyzed by counting the number of colonies. Graphs represent mean +/- SD. E) We seeded four million HCT116 cells per well. One day later a wound was created by scratching the cell layer vertically with a 100µL tip. Pictures were taken at this initial time, then cells were treated the same day and each day with 10mM DCA, 2ng/mL TGFβ-2 and/or 5µM of XMD8-92. Representative images of the wound healing at initial time and after 40h are shown. F) The area of the wounds of 3 independent experiments were calculated per condition with the software “ImageJ”. Graphs represent mean +/- SD. G-I) HCT116 cells were seeded at 2×10^5^ cells per well in 6 wells plates and treated for 24h with 10mM DCA or 2ng/mL TGFβ-2. RNA was extracted and mRNA gene levels quantified. These were split in metabolic genes (G; *CD36, ACDVL and ERK5*), EMT-related genes (H; *SNAIL, ZEB1, TWIST*) and NK cell ligands (I; *MICA, MICB, ULBP1 and ULBP4*). The data were analyzed with the 2^ (-DDCt) method and normalized to *β-actin* mRNA levels. The graphs represent mean+/-SD of the relative mRNA (n=3). * p < 0.05, ** p < 0.01, *** p < 0.001, **** p < 0.0001; Tukey’s test was used to compare condition to control or as depicted in the graphic.

ERK5 pathway is involved in EMT and cell migration(*35, 37*). To investigate the effects of DCA on cell migration, we performed transwell assays. DCA increased HCT116 migration and co-addition of OA did not increase DCA effects (Fig. 2C-D); although OA *per se* was able to induce migration. This could indicate that both compounds propel a similar metabolism. XMD8-92 completely blocked cell migration. In wound healing experiments, another model of metastatic potential, DCA induced wound closure similarly to TFG-β (Fig. 2E-F), a factor that induces EMT at least partially by promoting FA metabolism(*38*). XMD8-92 significantly inhibited wound healing (Fig. 2E-F) and reversed the effects of DCA. TGF-β induced the expression of *CD36, ACADVL* and *ERK5* along with EMT-related genes such as *SNAIL, ZEB1* and *TWIST* (Fig. 2H), but to a lesser extent than DCA (Fig. 2G-H). Finally, both TFG-β and DCA increased the expression of stress ligand genes (Fig. 2I), suggesting that changes in lipid metabolism may be implicated in multiple physiological processes (see below).

### DCA favors cell motility and invasion in 3D cultures

To further assess the impact of DCA on HCT116 cell migration, we generated 3D tumor spheroids in ultra-low attachment (ULA) plates. DCA significantly reduced the spheroid area after seven days of treatment (Fig. 3A-B), showing that DCA also had cytostatic activity in 3D tumor cell cultures, as in 2D (*27*). In addition, DCA significantly increased the number of detached cells around the spheroids (Fig. 3 A, C). We removed and pooled the spheroids, and separately pooled the remaining medium containing detached cells for each condition (Fig. 3D-E). DCA increased the expression of the same metabolic (CD36 and ACADVL) and EMT (ERK5, SNAIL and ZEB1) genes in 2D and 3D cultures. An increase was found in both the cells that stayed in the spheroid (Fig. 3D) and the cells that detached (Fig. 3E), with the exception of ACADVL, which did not change in detached cells. MHC-I was also increased in attached cells, but not in detached cells. We also found that DCA increased the expression of the activating ligand for NK cell *ULBP4* and *PVR* in HCT116 cells grown in spheroids (Fig 3D-E).

**Figure 3.**
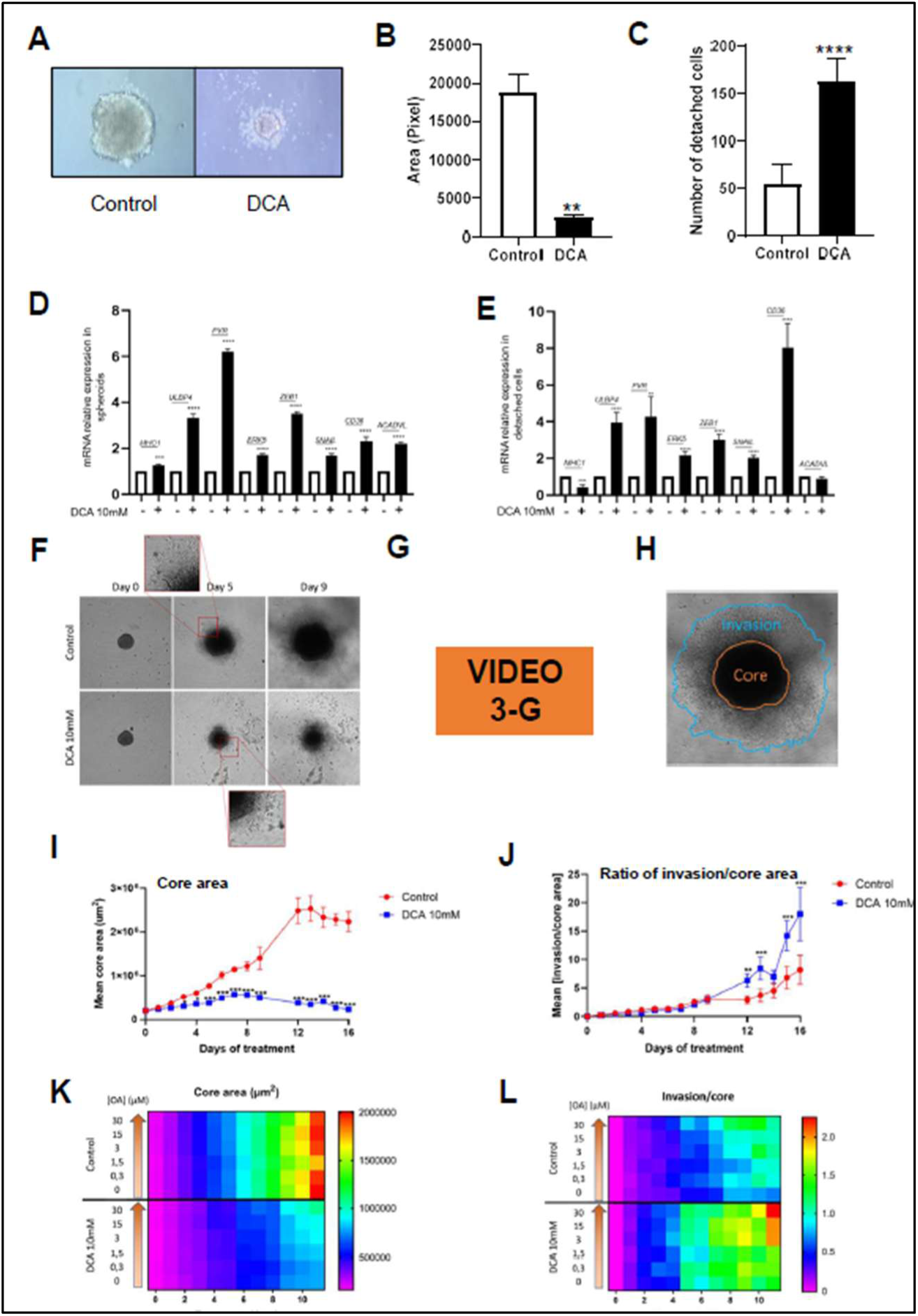
DCA decreases tumor growth and increases tumor cell detachment and invasion in *in vitro* 3D models. A-D) HCT116 were grown to generate spheroids in “Ultra Low attachment plates” in DMEM F12 B27 + Insulin/EGF/FGF medium and treated with 10mM DCA for 7 days. Images A) were taken, B) the area of the spheres and C) the number of cells detached from them were analyzed with the software “ImageJ”. The graphs represent the mean +/- SD. D-E) Spheroids were individually collected from the wells and pooled. The remaining detached cells were also pooled for each condition. 96 wells were pooled for each condition to obtain enough RNA, which was extracted and RT-QPCR performed for each condition: spheres (D) and detached cells (E). The data were analyzed with the 2^ (-DDCt) method and normalized to *β-actin* mRNA levels. The graphs represent mean+/-SD of the relative mRNA (n=3). * p < 0.05, ** p < 0.01, *** p < 0.001, **** p < 0.0001; Tukey’s test was used to compare condition to control or as depicted in the graphic. F-L) HCT116 cells were cultivated in ultra-low attachment 96-well plate. When spheroids were formed, they were transferred in U-bottom 96-well plate and resuspended in GeltrexTM. Treatments were performed every day from day 0 and during 16 days and images were taken on a Cytation 5. F) Representative pictures of spheroids growth and tumor invasive profile in GeltrexTM. G) Video of cell invasion at different time points, with DCA-treated cells at right of the images. H) Schematic representation of the sphere (the core) and the invasive area (in blue) generated by the cells around the sphere. I) Growth curve of the spheroids’ core. J) Invasiveness is measured by the ratio between the invasion area and the core area. K-L) Heat maps of the core (K) or the invasion area (L) of spheroids growing with or without 10mM DCA in the presence of different concentrations of OA. We analyzed 6 wells per condition and the experiment were replicated 3 times. All the results were treated by GraphPad Prism 9 software with multiple t test; statistical significance determined using the Holm-Sidak method; alpha = 0.05. * p<0.05; **p<0.01, ***p<0.001, ****p<0.0001. * p<0.05; **p<0.01, ***p<0.001, ****p<0.0001; unpaired T test.

Next, we analyzed the invasive capacity of the spheroids treated with DCA. HCT116 cells were cultivated in ULA 96-well plates and after spheroid formation, they were transferred onto a 7% Geltrex matrix and treated with 10mM DCA each day (Fig. 3F-J). Without treatment and after 13 days, the control spheroid reached an average 2.48×10^6^ µm^2^ whereas DCA treated spheres only 3.8×10^5^ µm^2^ (Fig. 3F, I). Thus, DCA limited spheroid size. Video capture shows the effects of DCA throughout the 16 days (Fig. 3G). In contrast, DCA favored cell migration and invasion of the surrounding environment and these invasive cells changed their phenotype (Fig. 3 F-H). With DCA, the cells detached from each other and from the spheroid. To overcome the inherent problem of the variable spheroid sizes, we quantified the area that was occupied by cells outside the spheroid core, and called this “invasion area” (Fig. 3H, J), then determined the ratio between the invasion area and the core area (Fig. 3J). DCA increased this ratio after 13 days in culture, indicating that it increased cell detachment and mobility in 3D structures. The addition of OA did not affect the core size of the spheres (Fig. 3K). However, it strongly increased the invasion/core ratio, especially in the presence of DCA (Fig. 3L), suggesting the involvement of lipid metabolism.

### FAO induces expression of stress ligands on tumor cells, recruitment of NK cells and sensitization to NK cells

We recently showed that DCA increases the expression of genes encoding NK cell-activating ligands in leukemic cells (*39*). As mentioned above, in 2D culture conditions, DCA also able to increased *MICA, MICB, ULBP1 and ULBP4* in HCT116 cells (Fig. 2I). *ICAM-1* expression also increased. Hence, FAO induces the expression of a pattern of ligands that should favor NK cell recognition and NK cell-mediated elimination. Furthermore, DCA induced stress ligand expression in both spheroids and detached cells (Fig. 3D-E). Hence, we analyzed whether DCA favored NK cell recognition by tumor cells. First, we incubated CTFR prestained *in vitro* expanded and activated NK cells (eNK) with HCT116 spheroids for 48h, harvested the spheroids and surrounding medium separately, and analyzed the cells by flow cytometry (Fig. 4A). DCA increased NK cell infiltration in spheroids (Fig. 4A1). As expected, a reduction in the non-infiltrate fraction was observed (Fig.4A2). To visualize NK cell infiltration, we stained HCT116 spheroids with calcein before adding the CTFR-stained eNK cells. DCA-treated spheroids showed a significant increase in NK cell infiltration (Fig. 4B-C). We also used automated microscopy Cytation®, which confirmed that DCA promoted eNK to deeply infiltrate the spheroids (Supplemental Fig. 1A). This was even more evident when spheroids formed from MDA-MB-231 cells (Supplemental Fig. 1B). As DCA recruited eNK towards the spheroids, we investigated if it could favor tumor cell destruction. In 2D, DCA did not significantly increase eNK cytotoxicity (Fig. 4D). However, in 3D DCA and eNK separately reduced spheroid area and when used together they significantly reduced it further (Fig. 4E). Notably, DCA did not reduce the surrounding (halo) area, which mainly comprised eNK and the remaining detached cells (Fig. 4F). Supplemental Fig. 2A shows that eNK cells moved to the center of the well in the absence of spheroids. To exclude the possibility that eNK moved to the spheroids due to gravity, we placed the spheroids out of the well center (Supplemental Fig. 2B). We observed that eNK cells moved to the spheroid even if certain cells remained in the center. This showed that NK cells infiltrated the spheroids, which was further enhanced upon DCA treatment.

**Figure 4.**
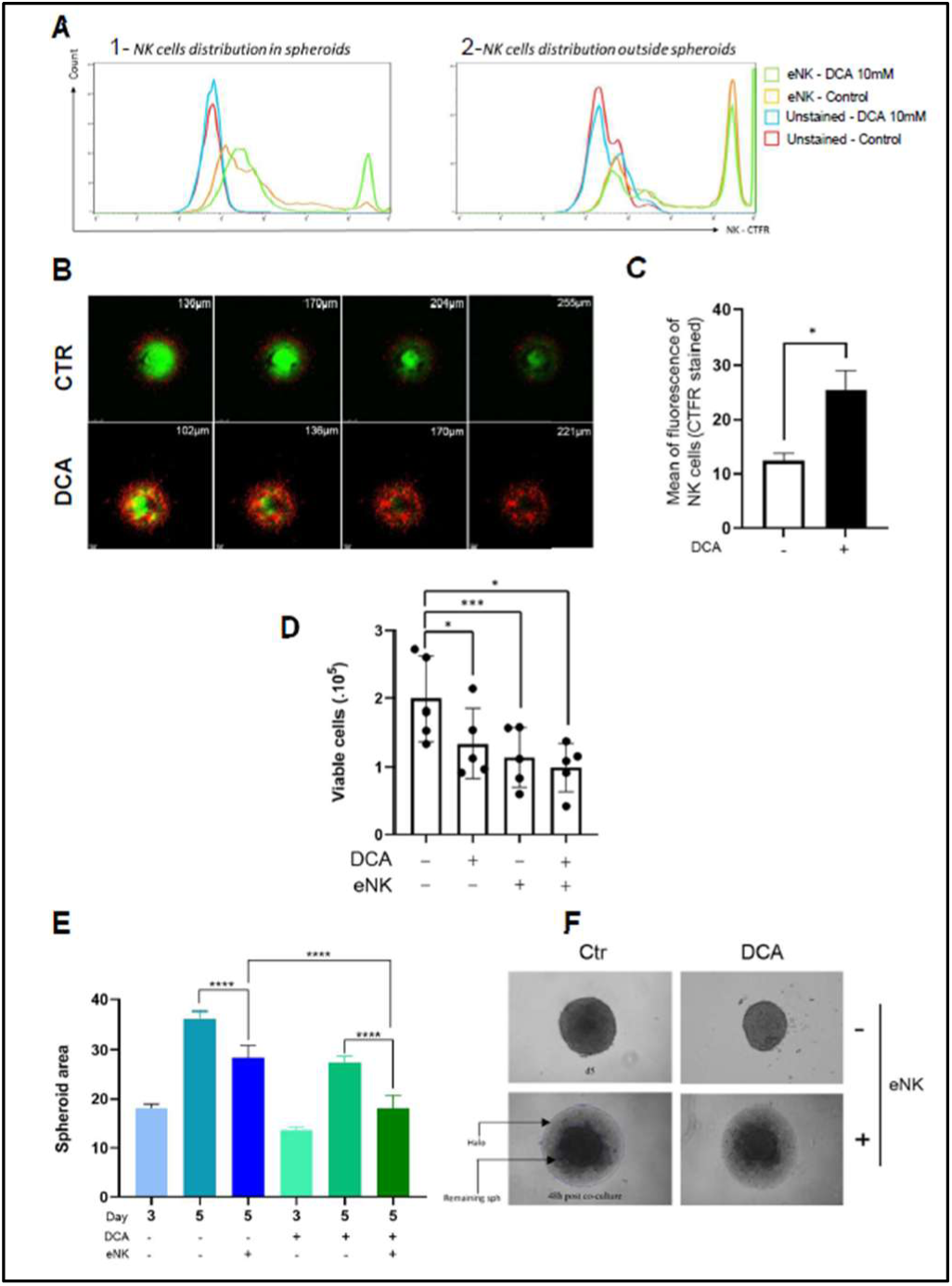
DCA increases NK cell infiltration into tumor cell spheroids and sensitizes tumor cells to NK cell-mediated killing. To assess eNK infiltration in spheroids both flow cytometry (FC) and confocal microscopy were used, A) FC: spheroids pre-treated with DCA or H2O (DCA vehicle) for 48 hours were cocultured with CTFR-stained eNK cells for 8 hours. We collected separately spheroids and the remaining cells in the media to compare the infiltrated and the media’s remaining fraction of eNK cells. Cell fluorescence was assessed: CTFR positive population in the right side of the graphs represents NK cells whereas spheroids tumor cells are the negative population. B) Confocal microscopy: Calcein-AM (green) stained spheroids and CTFR stained eNK cells (red) were co-cultured for 24h, and imaging was processed by confocal microscopy (Leica-SP8). Images represent different confocal planes of the spheroids. C) Fluorescence intensity was analyzed with the software “ImageJ”. The graphs represent the mean ± SD. D) 2.10^5^ HCT116 cells were plated in 24-well plates and treated with 10mM DCA or H2O for 24h then incubated with eNK at an effector: target (E:T) ratio of 2:1. After 6 hours, HCT116 cells were counted. Graphs show mean ± SD of 5 independent eNK expansions. Statistical analysis was performed by the paired t test using the GraphPad Prism 9. E) After seeding, tumor spheroids were treated daily for 3 days with 10 mM DCA or H2O. eNK cells were added at E:T ratio of 2:1 for 48h. Cytotoxicity (spheroid destruction) was monitored by brightfield microscopy, and Fiji software measured and analyzed the solid spheroid area as described in (F). The graph represents the mean ± SD of the sphere area of 24 spheroids which were processed in 2 independent days, i.e. 12 spheroids/experiment. Statistical significance was analyzed by one-way ANOVA followed by Tukey’s multiple comparison test. * p<0.05; **p<0.01, ***p<0.001, ****p<0.0001.

### FAO increases metastasis in a zebrafish model

We have shown in several *in vitro* models that DCA induces EMT in cancer cells, facilitating their migration and invasion. To assess this property in vivo, we chose zebrafish as a suitable model (*40*), in which we could investigate cell motility in individual cells. We used the MDA-MB-231 cell line because of its invasiveness and aggressiveness (*41*), and evaluated metastasis by counting the number of fish in which cells were observed in the zebrafish body away from the injection site (periyolk space), specifically in the tail, at various time points. Epifluorescence images obtained after in vivo monitoring at 24, 48, 72, and 120 hours post injection (hpi) allowed us to determine the effect of DCA on cell invasion properties (Fig. 5A). DCA-treated MDA-MB231 xenografted cells were observed in more than 80% of fish at 24hpi, compared to only 57% of untreated cells at the same time point (Fig. 5B). However, at 120hpi, we did not observe statistical differences, as the fraction of invasive cells in non-treated fish increased to reach 100% under both conditions (Fig. 5B). High-resolution images showed single-cell migration to the perivascular caudal region at 24hpi (Fig. 5C). In DCA-treated cells, invasive cells were more numerous at the periphery of the dorsal longitudinal anastomotic vessel (DLV) and posterior cardinal vein (PCV) in the zebrafish caudal region than in non-treated cells. Moreover, we noticed that DCA pretreated cells were more dispersed after injection than non-treated cells. To assess whether FAO enhances the number of migrating cells, we measured the area of the invading cells (Fig. 5D). At 24hpi, the mean area tended to increase in the DCA-treated condition compared to than in the non-treated condition, although this effect was lost at later time points.

**Figure 5.**
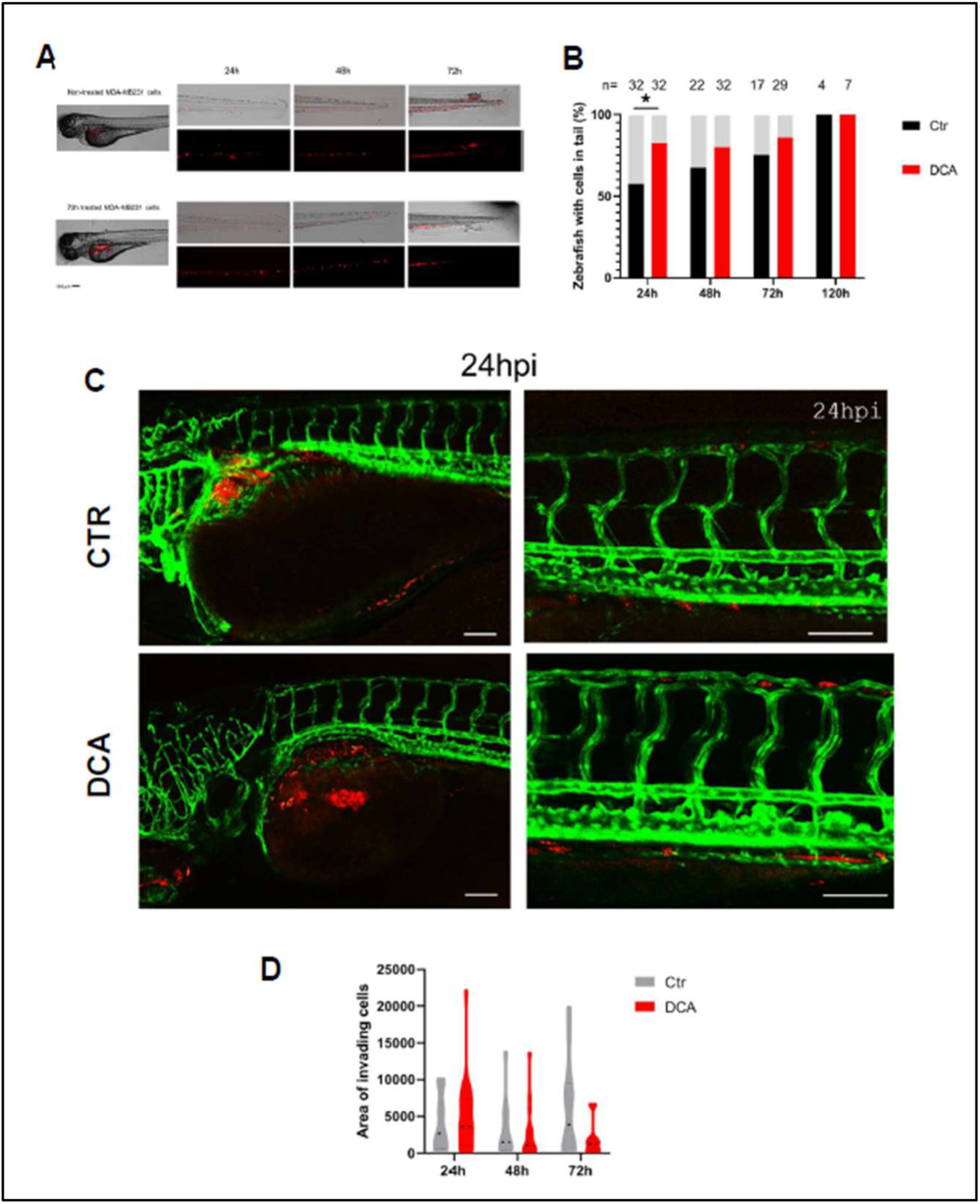
DCA induces invasion *in vivo* in a zebrafish model. A) In a zebrafish invasion assay, MDA-MB231 cells pretreated or not with 10mM DCA were labelled with CTFR and injected in the periyolk of 48hpf zebrafishes larvae. Epifluorescence images (inverted confocal microscope TCS SP5, Leica systems) were taken at 24, 48, 72 and 120h and cell migration was quantified. Scale bar represents 100 µm. B) The number of zebrafish with metastatic cells in the tail was counted at 24, 48, 72 and 120h post engraftment. Statistical differences were analyzed using chi square test, * p<0.05. C) DCA enhances perivascular metastasis in the caudal part at 24h post injection (hpi). The same human cells were equally engrafted in transgenic zebrafish line: Tg(Kdrl:egfp to visualize the vascular system (green). High-resolution images were acquired using a confocal microscope. Scale bar=100µm. D). The area of invading cells in zebrafish with metastasis was measured to assess if the DCA increases the number of metastatic cells.

### DCA favors metastasis formation in mice

To further confirm the ability of DCA to induce metastasis we engrafted 4T1 cells orthotopically into athymic nude fox n1 mouse mammary fat pads. We chose nude mice because they lack T and B cell compartments but contain NK cells(*42*) allowing the investigation of the effect of NK cells on metastatic spreading. We administered DCA by gavage in order to have a good bioavailability of the drug. DCA did not affect bodyweight or mouse behavior and had no impact on the tumor area or volumes (Fig. 6A-B). However, it significantly increased the number of metastases per lung (Fig. 6C) and tended to increase the number of metastases per lobe (Fig. 6D). Therefore, DCA increased cell motility and metastasis in the two *in vivo* models.

**Figure 6.**
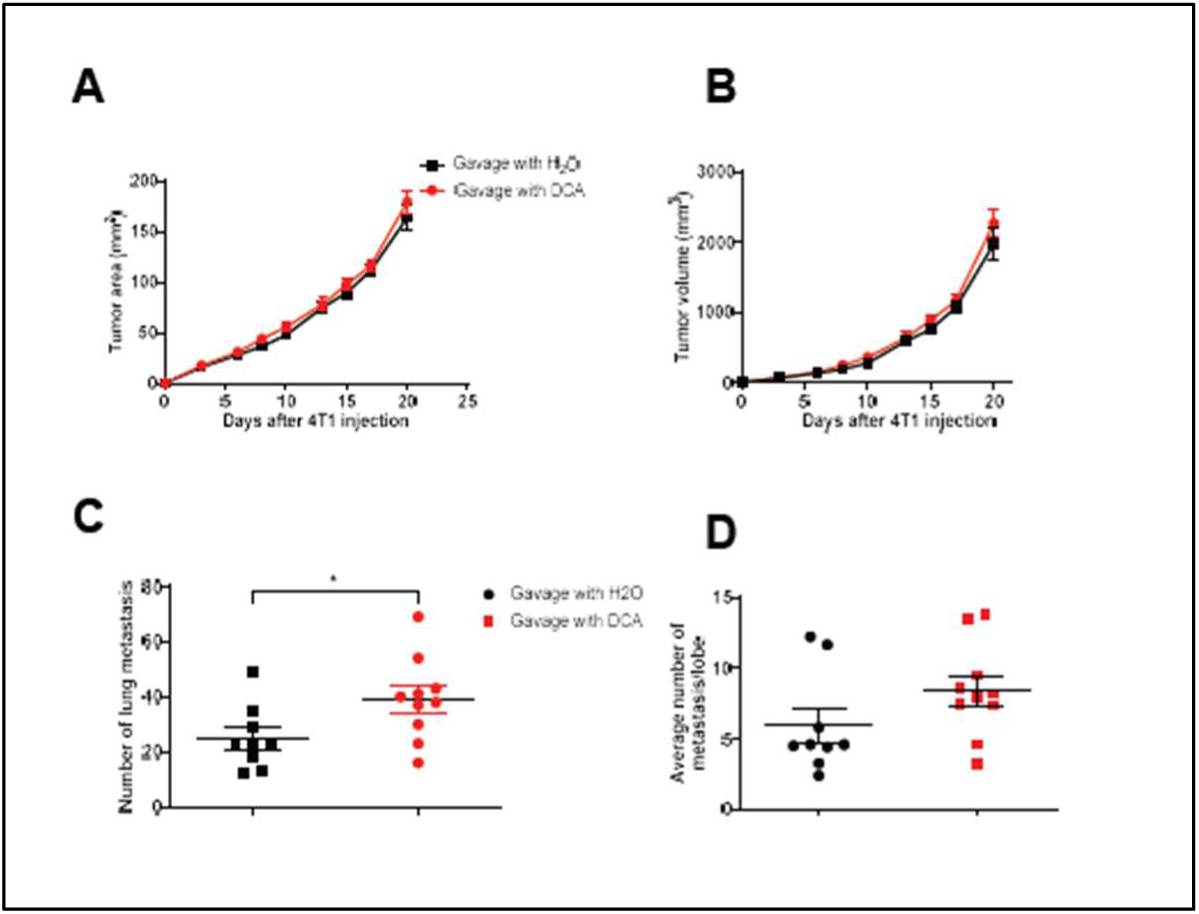
DCA favors lung metastasis. A) Tumor area (obtained by multiplying the length by the width of the tumor) after injection of 4T1 cells into Athymic nude mice, monitored for 3 weeks. Error bars, SEM (n = 10 mice). B) Tumor volume (obtained by multiplying the area by the size of the smallest axis). Error bars, SEM (n = 10 mice). C) Number of metastases in the whole lung. The different lobes of each lung were dissected and metastases were counted in each lobe. Horizontal bars represent the mean +/- SEM (n =10 for DCA and n = 9, for H_2_O treated mice). Unpaired t-test, two tailed p-value = 0.0347. D). After counting the number of metastasis in each lobe, the average number of metastasis per lobe was calculated and plotted. Horizontal bars represent the mean +/- SEM (n =10 for DCA and n = 9, for H_2_O treated mice). Unpaired t-test, two tailed p-value = 0.1329.

### FAO sensitized tumor cells to NK cell-induced killing in vivo

To investigate whether metabolic changes sensitized tumor cells to NK cell-mediated killing, we treated MDA-MB231 cells with DCA and engrafted them alongside eNK cells in zebrafish, as shown in Fig. 5 (Fig. 7A and Supplemental Fig. 3). We observed that DCA-treated tumor cells exhibited increased presence in the tail region. eNK migrated through the vascular system of the zebrafish larvae and successfully reached the tail (Supplemental Fig. 3). In the presence of eNK cells, there was a significant reduction in the number of tumor cells reaching the tail, effectively preventing tumor cell invasion in this area (Fig. 7A). To quantify this reduction, we measured the fluorescence intensity in the tail as a readout of tumor cell invasion (Supplemental Fig. 4). To compare the effect of NK cells on control cells and DCA-treated cells,the fluorescence intensity in non-eNK treated fish was set to 100 (Fig. 7B). This enabled a comparative analysis, revealing that eNK cells were more effective among DCA-treated MDA-MB231 cells.

**Figure 7.**
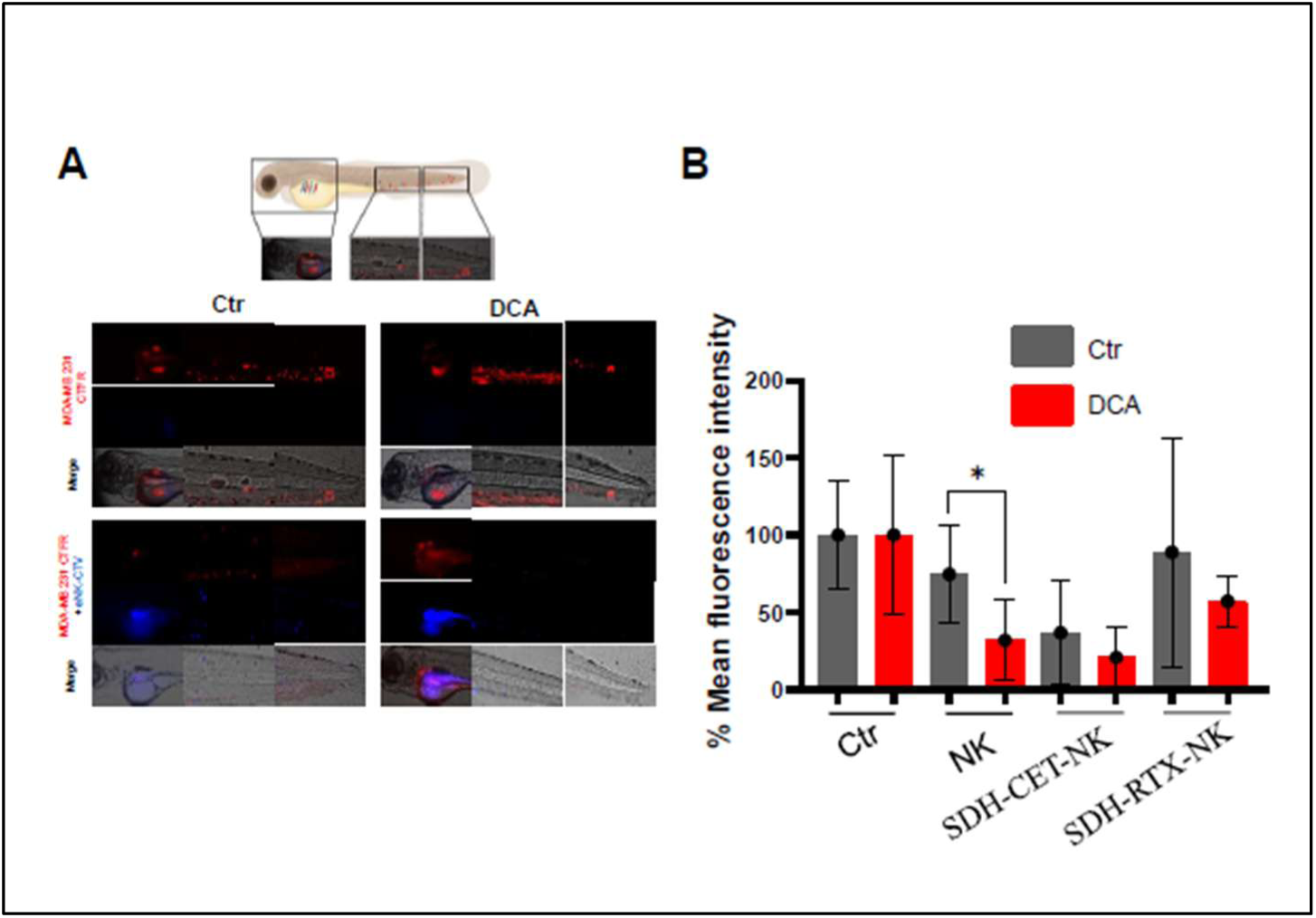
NK cells preferably target migratory cells. Epifluorescence images showing the transgenic line of zebrafish larvae tg(kdrl:GFP) (whole organism), images were taken at X10 magnification. A) MDA-MB231 cells pretreated or not with 10mM DCA were labelled with CTFR and injected in the periyolk of 48hpf tg(kdrl:GFP) zebrafishes embryo. eNK were armed, or not, with SDH-cetuximab and SDH-rituximab and labelled with cell trace violet and injected at the same time than MDA-MB231 cells with an E:T ratio of 5:1. Images (inverted confocal microscope TCS SP5, Leica systems) were taken 48h post injection (hpi) and the area of invading MDA-MB231 CTFR cells was quantified. Vessels are in green, MDA-MB231 in red, and eNK in blue. Scale bar = 100µm. B) Graph represents de mean+/-SD of three larvae of 2 independent experiments. The values of larvae non-engrafted with eNK were arbitrarily taken to 100 for both treated and non-treated MDA-MB231 cells and the percentage of fluorescence of the eNK-treated larvae was compared between treated and non-treated MDA-MB231 cells. Statistical differences were analyzed using chi square test, * p<0.05.

We recently demonstrated the ability of SDH-modified monoclonal antibodies (mAbs) to confer tumor antigen specificity to eNK cells by arming them via their CD16a receptors with SDH-modified monoclonal antibodies (mAbs) (*43*). To test this, we armed eNK cells with SDH-cetuximab, which targets EGFR, a receptor highly expressed in this triple-negative breast cancer cell line(*44*). As a control, we used SDH-rituximab, an anti-CD20 mAb, independent of this cell line. SDH-cetuximab-armed eNK showed greater efficacy than non-armed eNK, although differences were observed between the control and DCA-treated cells (Supplemental Fig. 3 and 4 and Fig. 7B). This may be due to the high effectiveness of SDH-cetuximab-armed eNK cells, which eliminated approximately 70% of the target cells (Fig. 7B). As expected, SDH-rituximab-armed eNK cells did not outperform the non-armed eNK cells. These results confirmed that DCA-induced metabolic rewiring sensitizes tumor cells to NK cell-mediated targeting.

## Discussion

FAO, which fuels β-oxidation and mitochondrial respiration, is associated with tumor progression, invasion, and metastasis (*5, 6*). To sustain FAO, metastatic cells increase FA import and activate various genes required for FAO, notably CD36 (*6, 45*). Our findings show that forcing FAO induces motility, invasion, and EMT-related gene expression, indicating a mutual enhancement between EMT and FAO. However, as metastatic cells gain motility and increase metabolic substrate availability, they become more susceptible to NK cell targeting. We propose that this originates from the same pathways that regulate FAO, which also induce the expression of tumor cell ligands recognized by NK cells, leading to tumor cell destruction. This phenomenon is observed primarily in 3D cultures and zebrafish models. In 2D settings, the effect is less significant, likely due to structural differences in NK-tumor cell interactions. In 2D, cells tend to spread rather than move; therefore, infiltration and/or motility are less essential for tumor recognition and/or killing. The inclusion of spheroids in Geltrex, even at 7%, better simulated a structured matrix, allowing cell motility that closely resembled physiology. Under these conditions, spheroids may secrete factors and cytokines that are absent or are less effective in 2D cultures(*46*), thereby enhancing NK cell recruitment and/or killing.

Consistent with previous studies in other cell types(*39*), FAO induced an activating NK cells ligand pattern in HCT116 cells. First, it increased *PVR* mRNA levels in both 2D and 3D cultures. PVR, a member of the nectin-like family, has emerged as a promising immunotherapy target for enhancing anti-tumor responses(*47, 48*). Similar to other family members, PVR plays a role in adhesion, contact inhibition, migration, proliferation, and immune response(*47, 48*). It is overexpressed in several cancers, promotes invasion, migration, and proliferation, and is associated with poor prognosis and aggressive tumor progression(*49*). PVR has substantial immunoregulatory potential, interacting with DNAM-1 or CD226 (*48, 50*), an immune-activating receptor, and with the inhibitory receptors T cell immunoglobulin and ITIM domain (TIGIT) and CD96 (*51*), balancing immune cell activation or inhibition, respectively. In healthy individuals, the balance between activating and inhibitory signals maintains normal immune function; however, in tumors, PVR overexpression often promotes immune escape. Targeting PVR can redirect the immune response to eliminate tumor cells (*51*). Chockley et al. demonstrated that increased PVR and decreased E-cadherin levels correlate with enhanced NK cell immunosurveillance and cytotoxicity (*11*). Second, FAO decreases the expression of two inhibitory tumor ligands, CDH1 (encoding E-Cad) and MHC-I. CDH1 interacts with KLRG1, and MHC-I with KIRs; thus, reducing E-Cad and MHC-I expression improves NK cell recognition. Third, in our 3D model, FAO also upregulated ULBP4, a stress ligand recognized by the NK cell-activating receptor NKG2D (*15*), which also binds other stress ligands, such as ULBP1 and MICA/B ^3^, which are elevated in our 2D models. NK cell activation depends on multiple receptors(*52, 53*), and the loss of inhibitory contact between E-Cad and KLRG1 alone does not reinitiate NK cell cytotoxicity(*11*). Thus, it is unlikely that any single ligand drives FAO-induced tumor cell sensitization to NK cells; rather, a pattern of ligands likely facilitates NK cell recognition. After DCA treatment, this ligand pattern may vary by disease and patient, potentially explaining the high variability in DCA response in clinical trials (*29, 54*). Currently, we have not fully identified this pattern and believe that it will be challenging to uncover.

Our zebrafish model showed transient effects of FAO on cell motility: DCA pretreated cells migrated more significantly at 24 hours, but this effect diminishes thereafter. Several factors could explain this observation. First, DCA is rapidly metabolized, and we hypothesize that after 24 h, the treated cells are no longer affected by DCA, losing their migratory advantage. Second, DCA decreased proliferation, potentially resulting in fewer cells after 24 h compared with the controls. Lastly, regardless of treatment, cells continue to “travel” in the host, with invasive cells reaching 100% in the fish away from the injection site by 72 h; therefore, differences are expected to diminish over time. In our mouse model, continuous DCA treatment increased the metastasis. Overall, we found that DCA enhanced motility and invasiveness in vivo, leading to increased metastasis.

Most cancer-related deaths are due to metastasis, for which there are currently no effective treatments(*55*). EMT enables epithelial cancer cells to adopt a mesenchymal phenotype, allowing them to detach from the primary tumor. ERK5 activation promotes EMT and is a marker of poor prognosis, for example, in triple-negative breast cancer and endocrine-resistant breast cancers(*56–58*). Pharmacological inhibition(*59*) or genetic ablation(*60*) of ERK5 can reverse EMT and the migratory phenotype, suggesting that ERK5 inhibition can treat patients with or at risk for metastasis. Targeting ERK5 effectors involved in FAO, FA synthesis, and/or uptake(*10, 61*), such as CD36, may be beneficial. This is reinforced by several facts: the overexpression of CD36 promotes metastasis(*6, 62*), and is linked to poor prognosis across several cancer types(*6, 7, 63*).

Athymic nude mice possess NK cells (*42*), and we observed an increase in metastasis in DCA-treated nude mice. This suggests that mouse NK cells were unable to control the spread of metastatic cells in our model. However, 4T1 cells are highly aggressive and spontaneously metastasize to multiple organs(*64*), making them difficult to control using endogenous NK cells. Notably, human and mouse NK cells exhibit significant differences in development, location, and physiology. Therefore, it is not possible to determine whether DCA promotes metastasis in humans. Although DCA has not consistently demonstrated anti-tumor clinical efficacy(*54*), it remains of major clinical interest(*29, 30*). Our laboratory previously showed that DCA upregulates cholesterol and lipid metabolism via the ERK5 pathway (*31, 32*). Here, we investigated the consequences of this activation. DCA demonstrates a dual effect; it significantly reduces spheroid size while promoting EMT and migration, which could negatively impact the prognosis of DCA-treated patients. On the positive side, DCA enhances NK cell-mediated tumor cell sensitization, although in our mouse model NK cells did not control the increase in metastasis (see above). However, the primary focus of this study was to examine how metabolic rewiring of metastatic cells affects immune cell recognition. Therefore, we used relatively high DCA concentrations, exceeding those used in clinical settings, any relationship between our findings and DCA clinical effects should be considered cautiously. Available data on the effect of DCA on metastasis in animal models are limited and mainly involve DCA in combination with anti-cancer drugs, generally showing that DCA reduces metastasis(*65–67*), likely due to its cytostatic and NK cell-activating roles. DCA has not been clinically indicated, likely due to its neurotoxic effects(*29, 30*), and our study highlights the potential side effects that physicians should consider. Discoveries in cancer initiation and progression pathways have led to the development of specific tumor inhibitors(*68*), while immunotherapies(*69*), including NK cell-mediated cell therapy(*53*), have also advanced personalized, specific treatments. NK cells are promising candidates in this field, but their clinical efficacy requires futher improvement. The development of drugs that modify NK cell tumor recognition could advance NK cell-based therapies(*22, 70*). The findings and tools from this study may support the discovery of new drugs with similar DCA effects and anti-metastatic activity. For instance, metformin, which sensitizes tumor cells to NK cells, may be a candidate(*26*). In conclusion, during EMT and metastatis, tumor cells perform FAO partly via ERK5, which simultaneously induces a pattern of ligands recognizable by NK cells (Figure 8), presenting an Achilles’ heel for metastasis in new environments.

**Figure 8.**
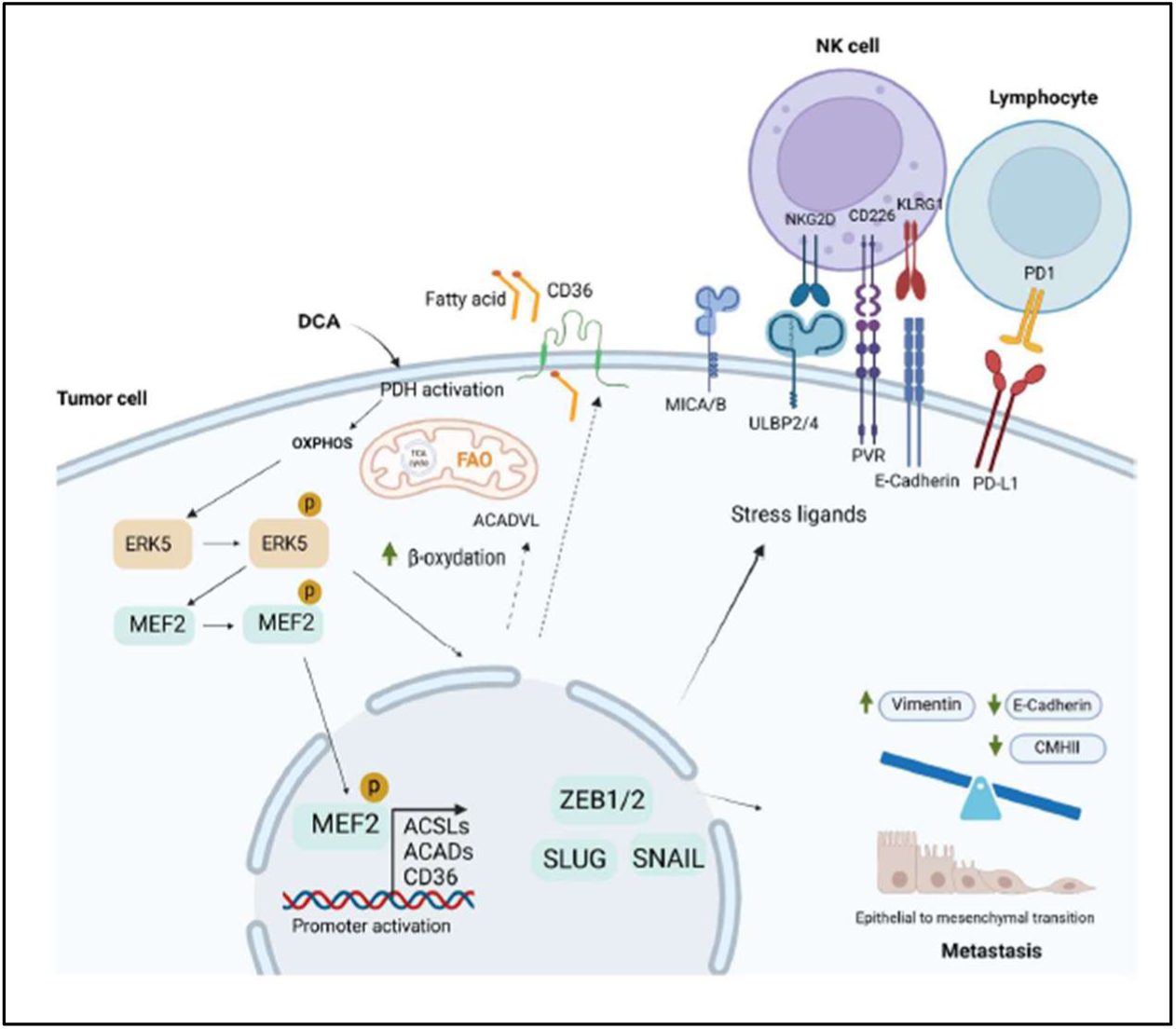
The relationship between lipid metabolism, metastasis and NK cell-mediated tumor immune surveillance. Forcing tumor cells to perform fatty acid oxidation activates the ERK5 pathway, which induces expression of proteins involved in lipid metabolism such as CD36, ACSLs and ACADs ^76^. In addition, it also induces different transcription factors implicated in the epithelial to mesenchymal transition including ZEB1/2, SLUG, SNAIL, which facilitates a metastatic phenotype. This changes in metastatic cells facilitates expression of stress ligand such as MICA/B, ULBP2, and PVR increasing NK cells recognition by the activating receptor NKG2D and CD226. Changes in other proteins such as downregulation of E-Cad, a ligand of the inhibitory receptor KLRG1 could also favor NK cell recognition.

## Methods

### Cell culture and transient transfections

The colon carcinoma HCT116 and the triple negative breast cancer MDA-MB231 cell lines were cultivated in 75cm2 flasks (Corning) in DMEM Glutamax containing Glucose (4.5g/L), pyruvate (0.11g/L) and 10% Fetal Bovine Serum (without antibiotics). Cell number, viability and death were analyzed using a Muse Cell Analyzer (Millipore) by incubating cells with Muse Count & Viability and Annexin V and Dead Cell kits respectively, following the manufacturer’s instructions.

For cell transfection, 2 million HCT116 cells were plated, and after 48 h they reached 80% confluency. Then, 300 μl of Opti-MEM (Invitrogen) was taken separately in two 5ml polypropylene tubes for each condition and 40 μl of Lipofectamine 2000 transfection reagent RNAiMAX (Invitrogen) was added to one of them and 12 μl of siRNA (20μM) was added to another tube for each condition following the manufacturer’s protocol. All siRNAs, that is siCD36, siERK5 and siControl were ON-TARGETplus SMARTpools (mixture of 4 siRNA) from Dharmacon. The cells were then analyzed after 48 h.

### Cell Migration Assay

Cells (25×10^3^ in 200 µl of medium without FBS) were seeded in the top chamber (insert) of 24-well transwell plates (Corning), and treated with 10mM DCA and/or 3µM OA. The outer chambers contained 700μl of the same conditioned medium with 10% FBS, except for the negative control, which did not contain FBS. This treatment was repeated for 24 h. At 48h, the medium was removed from both the insert and well. The inserts were washed twice with 1 x PBS, and cells were fixed in 4% paraformaldehyde (PFA) for 10 min. The cells were stained with crystal violet (24 h) and washed twice with 1X PBS. Staining from the upper layer of the well was removed using a cotton-tipped applicator and the inserts were allowed to dry. Finally, the lower layer (cells that reached the underside of the insert) was analyzed under a microscope and pictures were taken before analyzing the number of colonies.

### RT-QPCR experiments

Total RNA was extracted using NucleoSpin RNA isolation columns (Macherey-Nagel), reverse transcription was carried out using the iScript^TM^ cDNA Synthesis Kit (Bio-Rad) and 100μL of RNAse free water was added to the 20μl sample. Two µl of each sample were analyzed in a light cycler 480 Roche. All samples were analyzed using the 2 (-DDCt) method(*71*) and normalized to *β-actin* mRNA levels.

### Mito stress test (Seahorse XF)

Agilent Seahorse XF96 Cell Culture Microplates were pre-coated with poly-d-lysine. Then 7,5×10^3^ HCT116 cells were seeded in regular growth medium and treated with 5mM DCA and/or 3µM OA. The medium was changed after 24h and analyzed at 48h. Thirty minutes before the analysis, the seahorse was calibrated for the Mito stress test. The microplates were loaded and analyzed for 90 min. The oxygen consumption rate was determined using an XFe/XF analyzer.

### Immunoblotting

Samples were collected, washed with PBS and lysed with RIPA buffer. Protein concentration was determined by BCA assay (Pierce) before denaturation and electrophoresis on 4–15% TGX gels (BioRad). Protein transfer was performed using a TransTurbo system (Bio-Rad) on nitrocellulose membranes (Bio-Rad). After blocking for 1 h with 5% non-fat milk, the membranes were incubated overnight at 4 °C in agitation with primary antibodies, washed three times with PBS-Tween 0.1% and incubated with the appropriate HRP-labeled secondary antibody for 1 h. Membranes were washed out three times with PBS-Tween 0.1% and developed with Substrat HRP Immobilon Western (Millipore). Band quantification was performed using the “ImageLab” software from BioRad and represented as the ratio between the protein of interest and a control protein i.e. actin or tubulin. The value of 1 is arbitrarily given to control cells. Anti-CD36 antibody was from Gentex (GTX112891). EMT related protein antibodies were from the EMT antibody sampler kit (Cell Signaling, #9782). The antibody against β-Actin and HRP-labeled secondary antibodies were from Sigma.

### Immunofluorescence

Thirty-thousand HCT116 cells were seeded on coverslips into 6-well plates and treated for 48h with sterile H_2_O water (vehicle) or 5 mM DCA and/or 3µM OA. On the third day, the medium was removed and each well was washed with PBS. Then the cells were fixed in 4% PFA for 15 minutes at RT. After fixation, cells were incubated at RT for 15 minutes with an anti-CD36 antibody (CD36-PE clone REA760, Miltenyi; 1/100) and washed twice with PBS. Then, they were incubated for 15 minutes with DAPI (1/1000) and washed twice with PBS. After mounting the slides, they were dried in the dark for at least 2 days at 4°C and analyzed with a Leica Sp8 microscope.

### Neutral lipids levels (Bodipy)

It was basically performed as previously described (*39*). For immunofluorescence analysis, HCT116 were seeded on poly-D-lysine coated coverslips at day 0 and treated with/without 5mM DCA and 3µM OA for 48h. Cells were fixed with 4% PFA, and stained with Bodipy (2µM, 15min at 37°C) and DAPI. Images were taken with a Sp8 Leica microscope.

For lipid quantification by flow cytometry, HCT116 cells were seeded at 1×10^6^ cells in 10 cm dishes and treated with 5mM DCA and 30µM OA for 48h. Cells were incubated with Bodipy for 15 minutes, wash with PBS and harvested after trypsinization. After centrifugation, cells were resuspended in PBS and analyzed in a Gallios Flow cytometer (Beckman). Results were analyzed with Kaluza Software.

### Flow Cytometry

Briefly, 1 × 10^6^ cells were stained with the appropriate antibodies in PBS, 2% FBS and incubated at 4°C for 20 min. Cells were then washed and suspended in 200–250 μl of PBS and 2% FBS and the staining was analyzed using a Gallios flow cytometer (Beckman) and its Kaluza software.

### NK cell expansion and culture (eNK)

NK cells were expanded from patient umbilical cord blood according to the protocol developed by our team (*72, 73*) and then cultivated in RPMI 1640 GlutaMAX™ + 10% FBS, at 37°C with 5% of CO2 until use. eNK cells were counted with the Muse® Cell Analyzer.

### eNK labeling

eNK cells were labeled for 20min with CellTrace^TM^ Far Red in RPMI 1640 GlutaMAX^TM^ without serum, washed with RPMI 1640 GlutaMAX^TM^ + 10% FBS and suspended in complete spheroid medium. They were seeded with the spheroids at a ratio effector:target of 1:1 and incubated at 37°C with 5% of CO_2_ in the AgilentBioTek BioSpa 8 to begin the imaging kinetic.

### eNK cell infiltration by flow cytometry

One thousand HCT116 cells were resuspended in serum-free medium containing 1μM Calcein-AM (Invitrogen®, ref. C1430) for 4 h at 37°C. Spheroids were formed into U-plate ultra-low attachment cluster 96 well round bottom plates (Corning® Costar®, 7007) and grown for 2 days, then treated with 10mM DCA for 2 additional days. On day 5, NK cells stained with CellTrace™ FarRed (Invitrogen®, ref. C34564, 1:3000 dilution) following the provider’s instructions and added to the HCT116 spheroids at E:T ratio 1:1. Next day, the spheroids and their supernatants were pooled together. After sedimentation of the spheroids, the supernatants are collected separately. Both were prepared for flow cytometry analysis.

### Spheroid formation

Four hundred HCT116 or MDA MB 231 cells were seeded in U-plate ultra-low attachment cluster 96 well round bottom (Costar, #7007) plates in 200µL of DMEM/F12 media supplemented with B27 (Invitrogen), 20ng/mL EGF (Peprotech), 10ng/mL β-FGF (Peprotech) and 5µg/mL of human insulin (Peprotech). Plates were then centrifuged twice (opposite sides) without brakes at 400g. Spheroid formation was followed daily under a Zeiss microscope.

### Spheroids in Geltrex^TM^

Spheroids were formed in ULA and transferred into a Falcon^®^ 96-well Clear Round Bottom plate (one per well) by pipetting 20µL of medium containing the spheroid with P200 tips. Geltrex^TM^ was diluted in cold PBS to obtain a 7% solution and 100µL were added on the transferred spheroids to resuspend them. The plate was centrifuged for 5min at 4°C, 300g, to eliminate bubbles. Then it was incubated at 37°C in 5% CO2 for 30-45min to polymerize the gel. Once Geltrex^TM^ polymerized, 100µL of complete spheroid medium was added, directly on the gel, with the different treatments: H_2_O (control), 3µM OA and/or 10mM DCA (dissolve in H_2_O). Treated spheroids were incubated at 37°C with 5% CO_2_ and treatments were renewed every day and invasiveness measured as previously described (*74*).

### Spheroid Imaging

The 96-well plate containing spheroids in GeltrexTM 7% were imaged daily with the Agilent BioTek Cytation 5. eNK co-culture imaging kinetic was realized automatically by the association of the BioSpa and the Cytation 5. The plate was transferred between the tools by a robotic arm every 2 hours.

### Measures of the spheroid core and invasive area

The measures of the spheroid core and invasive area (Example Fig.5C) were performed on the Agilent BioTek Gen5 software and reported on an Excel spreadsheet. The invasiveness was calculated by the ratio between the spheroid area and the invasive area.

### Mouse experiment

Mouse breast cancer 4T1 cells, tested for absence of pathogens with Envigo NSP I test, were grown at 60% confluency. One million cells in 100 µl of PBS were injected in the mammary gland of 7 weeks old athymic nude fox n1 mice. 20 mice (females) were used in the experiment; 10 of which were treated daily (from Monday to Friday) during 4 weeks with 100 mg/kg of DCA (gavage with 100 µl of a 20mg/ml solution of DCA dissolved in H_2_O) and the other 10 (control) were treated by gavage with 100 µl of H_2_O. Mice were weighed every 2 days and tumor volume was measured using a Caliper every 2 days. After 4 weeks of treatment, mice were sacrificed by cervical dislocation and tumors were extracted and measured. Lungs were injected with a needle through the trachea with a solution of 15% Indian ink in distilled water, until the lungs were completely swollen. Black swollen lungs were then extracted from the mice and fixed using Fekete’s solution (40 ml of glacial acetic acid, 80 ml of 37% Formalin, 580 ml of (100%) ethanol and 200 ml of water). Metastases in the lung were observed as white (non-coloured) dots and the different lobes were dissociated to accurately count the number of metastases in each lobe.

### Zebrafish in-vivo assay

For *in vivo* metastasis assay, MDA-MB231 cells were treated for 72h with 10mM DCA and labeled with 5µM CTFR (Thermofisher). Four hundred cells were injected in the perivitelline space of 48hpf (hours post-fertilization) zebrafish embryp after being anesthetized with MS-222 (200 mg/l, Sigma-Aldrich) and maintained at 33°C. A total of 32 fish were used for both conditions (n=32). DCA effects were assessed at 24, 48, 72 and 120hpi (hours post-injection) using epifluorescence microscopy (inverted confocal microscope TCS SP5, Leica systems). The area of invading cells was measured using Fiji software (*40, 75*).

For high-resolution imaging, 400 CTFR-labeled MDA-MB231 cells were injected into the perivitelline space of Tg(Kdrl:egfp) transgenic zebrafish line with fluorescent vasculature. At 48hpf zebrafish embryos were placed on glass-bottom petri dishes and covered with methylcellulose containing 0.003% tricaine (Sigma-Aldrich). Images were acquired using a Leica SP8 confocal microscope (Leica Microsystems).

### Cytotoxic assay of eNK cells in zebrafish

We injected eNK alongside the cancerous MDA-MB231 cell line into the perivitelline space of 48hpf zebrafish embryos. MDA-MB231 cells were pre-treated with DCA for 72 hours prior and then labeled with CTFR and suspended at a concentration of 80 × 10^6^ cells/mL to inject 4nL in embryos (200 cells/embryo). For the SDH-mAb arming, eNK cells were armed with SDH-cetuximab or SDH-rituximab by incubating them with 10 µg/mL of these mAbs for one hour at 37°C with 5% CO_2_ as previously described (*43*). The eNK were labeled with Violet Cell Trace and injected at an effector-to-target ratio of 5:1 into the perivitelline space of 48hpf embryos. Imaging was performed using an epifluorescence microscopy. The program Fiji was used to measure the mean fluorescence intensity of cells in the tail of the zebrafish embryos at 48hpi.

### Statistical analysis

All data were analyzed with the GraphPad Prism 9 software with various tests specified in the figure legends.

## Acknowledgements

We acknowledge the imaging facility MRI, member of the national infrastructure France-BioImaging supported by the French National Research Agency (ANR-/10-INBS-04, «Investments for the future») for the access to the cytometry platform and Gallios 3 Lasers cytometer. We acknowledge the tissue engineering facility CARTIGEN supported by the “FEDER/Region Occitanie” program, CHU Montpellier and the University of Montpellier for providing access to the Symphony A3 Cytometer. Umbilical cord blood units (UCBs) were obtained from the Biological Resource Center Collection of the University Hospital of Montpellier - (BIOBANQUES Identifier - BB-0033-00031, CHU Montpellier),

## Author contributions

S.Z., M.V. and D.G. participated in the research design. S.Z., S.E., B.H., A.B., C.A., C.J., N.D., A.S., J.P., M.D.T., and D.G. conducted the experiments. M.C., L.C., and N.A-V. provided fresh eNK cells. All authors contributed to data analysis, discussions and interpretation. G.B., A.A., M.V and D.G. supervised the study and drafted the paper. All authors read and approved the paper.

