## Supplemental Figures for "Metabolic rewiring of cancer cells induces metastasis via ERK5 but triggers recognition by NK cells"

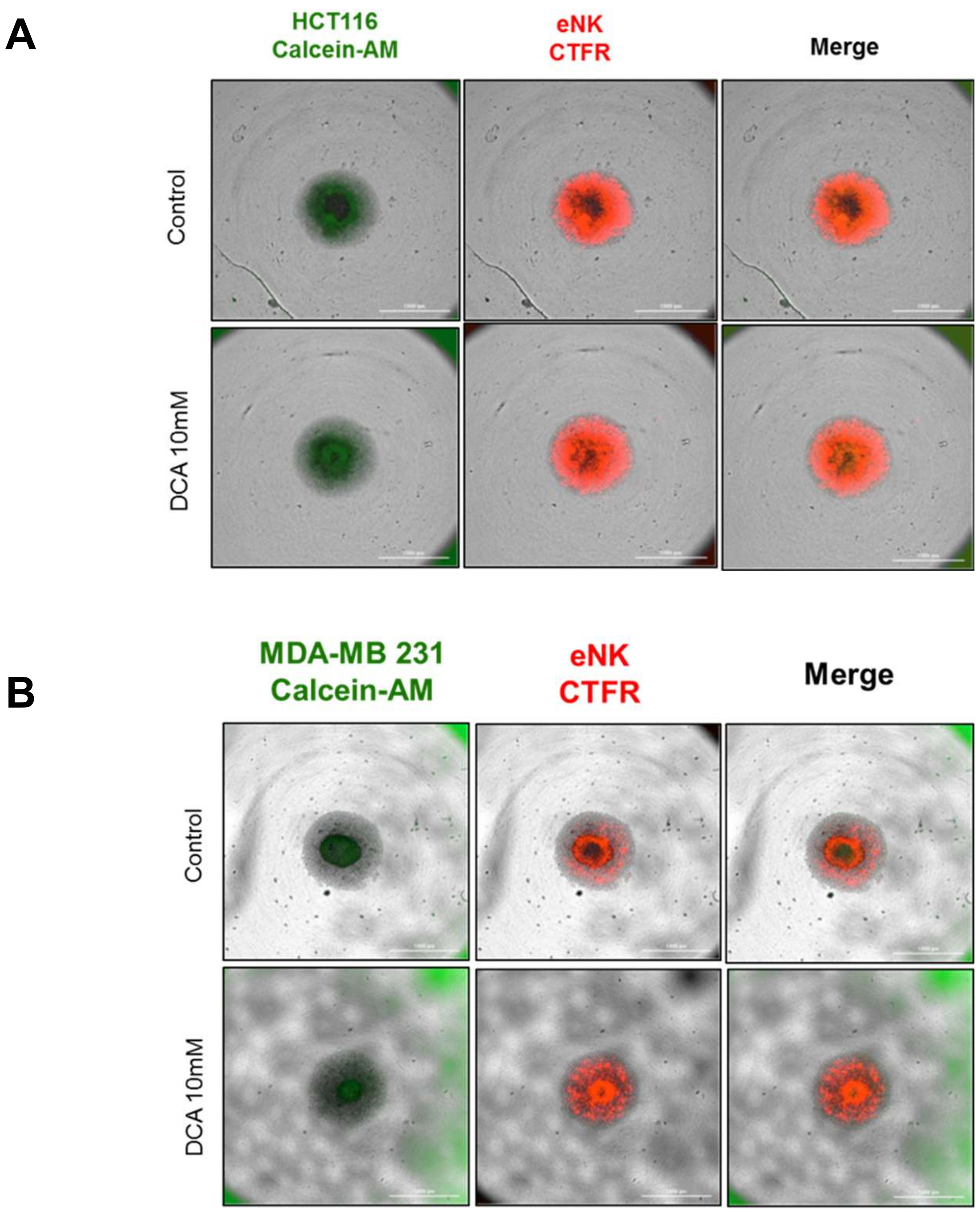


**Supplemental Figure 1. DCA increases eNK cell migration into spheroids.** HCT116 and MDA-MB 231 spheroids were stained with Calcein-AM (green) for 2h, and CFTR prestained eNK cells (red) were added for co-culture. Cellular imaging was processed with the Cytation5 automated microscopy 24h after eNK addition. Images are representative of 2 independent experiments with 3 spheroids per experiment.

### SUPPLEMENTAL FIGURE 1

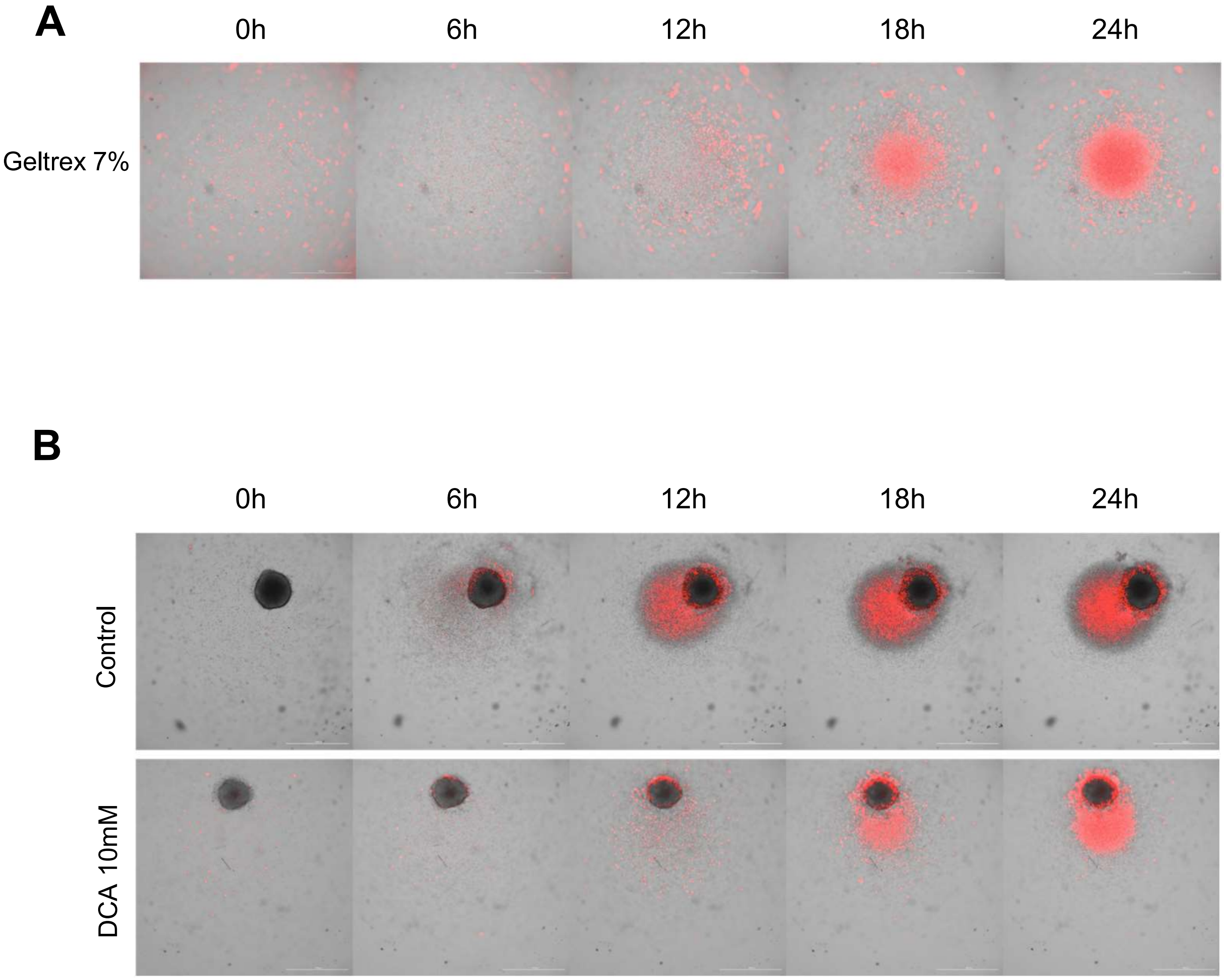


**Supplemental Figure 2. Migration of NK cells towards spheroids.** Before co-culture, eNK cells were labelled with CTFR. Labelled eNK were co-culture for 24h with spheroids included in Geltrex$^{TM}$ and treated with or without 10mM DCA during 4 days before eNK cells addition. Images were taken on a Cytation5. (A) Representative pictures of eNK infiltration in GeltrexTM in absence of spheroids (n=6). (B) Representative pictures of eNK and spheroids co-culture in GeltrexTM (n=6). The spheroids were voluntarily not placed at the center of the well to demonstrate specific movements of the eNK cells around the spheres.

### SUPPLEMENTAL FIGURE 2

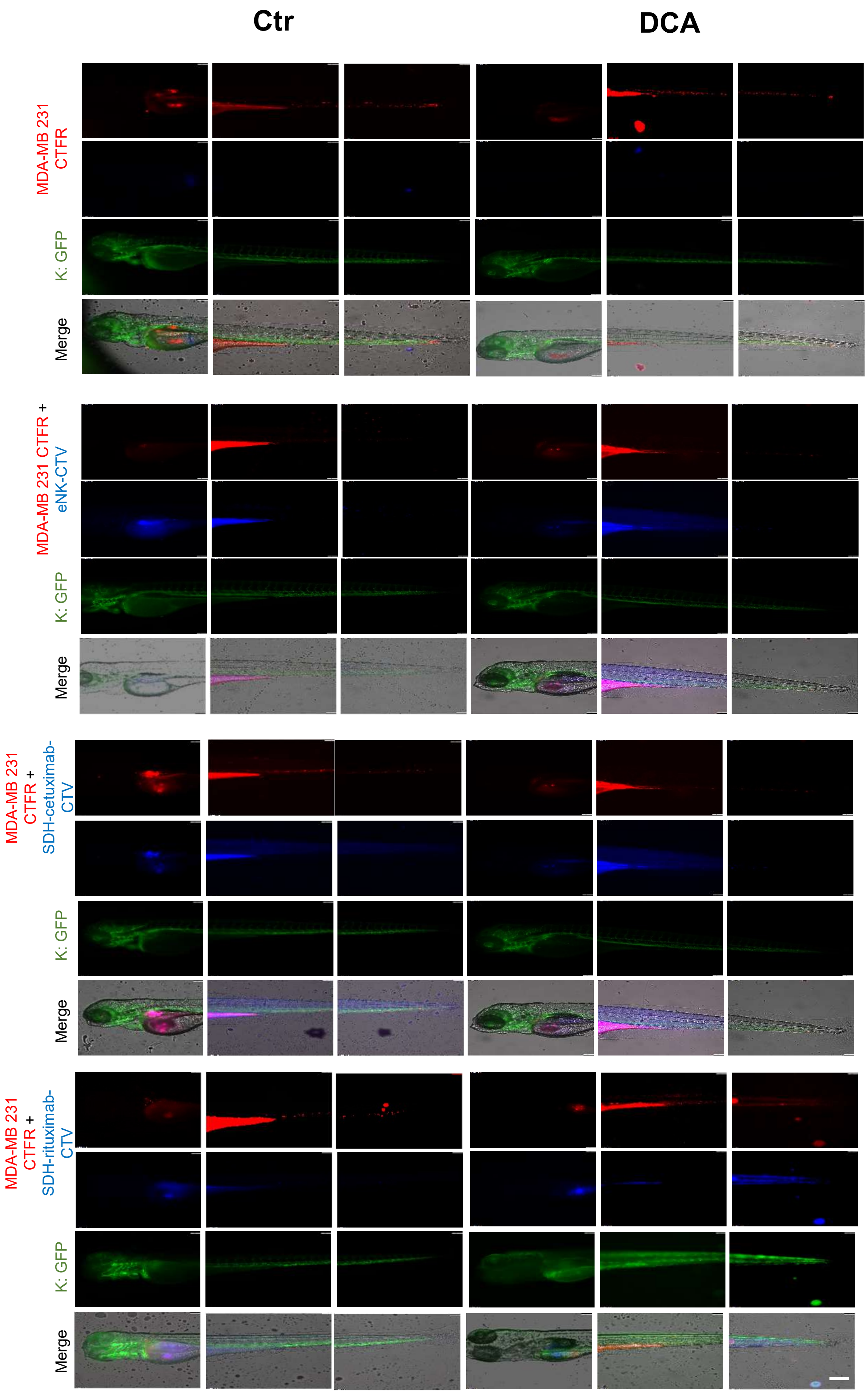


### SUPPLEMENTAL FIGURE 3

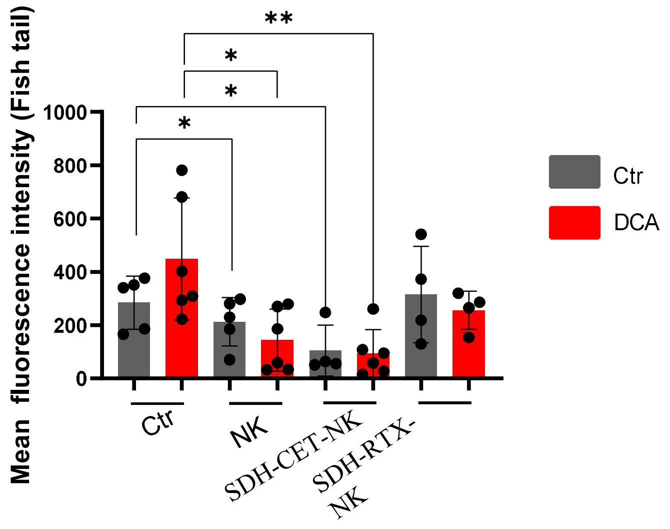


### SUPPLEMENTAL FIGURE 4
